# Metabolic snapshot of plasma samples reveals new pathways implicated in SARS-CoV-2 pathogenesis

**DOI:** 10.1101/2021.09.29.462326

**Authors:** Oihane E. Albóniga, Daniel Jimйnez, Matilde Sбnchez-Conde, Pilar Vizcarra, Raquel Ron, Sabina Herrera, Javier Martнnez-Sanz, Elena Moreno, Santiago Moreno, Coral Barbas, Sergio Serrano-Villar

## Abstract

Despite of the scientific and human efforts to understand COVID-19, there are questions still unanswered. Variations in the metabolic reaction to SARS-CoV-2 infection could explain the striking differences in the susceptibility to infection and the risk of severe disease. Here, we used untargeted metabolomics to examine novel metabolic pathways related to SARS-CoV-2 susceptibility and COVID-19 clinical severity using capillary electrophoresis coupled to a time-of-flight mass spectrometer (CE-TOF-MS) in plasma samples. We included 27 patients with confirmed COVID-19 early after symptom onset who were prospectively followed and 29 healthcare workers heavily exposed to SARS-CoV-2 but with low susceptibility to infection (‘nonsusceptible’). We found that the metabolite profile was predictive of the study group. We identified a total of 55 metabolites as biomarkers of SARS-CoV-2 susceptibility or COVID-19 clinical severity. We report the discovery of new plasma biomarkers for COVID-19 that provide mechanistic explanations for the clinical consequences of SARS-CoV-2, including mitochondrial and liver dysfunction as a consequence of hypoxemia (citrulline, citrate, and BAIBA), energy production and amino acid catabolism (L-glycine, L-alanine, L-serine, L-proline, L-aspartic acid and L-histidine), endothelial dysfunction and thrombosis (citrulline, L-ADMA, 2-AB, and Neu5Ac), and we found interconnections between these pathways. In summary, in this first report of the metabolomic profile of individuals with severe COVID-19 and SARS-CoV-2 susceptibility by CE-MS, we define several metabolic pathways implicated in SARS-CoV-2 susceptibility and COVID-19 clinical progression that could be developed as biomarkers of COVID-19.

## Introduction

Despite the effective response to the worst pandemic that humanity has faced in recent decades, the metabolic and biochemical processes during SARS-CoV-2 infection remain poorly understood. Most studies that have thus far investigated the biochemical pathways affected by SARS-CoV-2 rely on powerful bioanalytical techniques. Using untargeted and targeted metabolomics, other groups have identified that disruption of lipid and amino acid metabolism, such as the kynurenine pathway, are potentially relevant pathways associated with COVID-19 pathogenesis (1–5). Other candidate pathways that could be involved in clinical progression include pyrimidine (1,2) and purine (1,6–8) metabolism, fructose, and mannose metabolism (1,7) and carbon metabolism (1,2,9), although the specific mechanism remains unclear. Overall, the necessity to elucidate the global snapshot of biochemical processes behind SARS-CoV-2 infection is still in progress.

Metabolomic profiling can be performed by mass spectrometry (MS) coupled to a separation technique such as liquid chromatography (LC-MS), gas chromatography (GC-MS) or capillary electrophoresis (CE-MS). CE-MS is used to study polar and ionizable compounds such as free modified amino acids (MAAs) and “epimetabolites”, which are side products of enzyme reactions. These MAAs or the appearance of epimetabolites has been associated with important alterations in cellular, physiological, and pathological processes(10–13). While CE-MS is a powerful method to characterize unknown mechanisms of disease progression, to our knowledge, it has not been used in individuals with COVID-19.

Here, we investigated novel metabolic pathways of SARS-CoV-2 susceptibility and COVID-19 clinical progression using CE-MS in longitudinal plasma samples from patients with COVID-19 with different disease severities and in a population of healthcare workers heavily exposed to SARS-CoV-2 but with low susceptibility to infection.

## Results

### General characteristics of the study population

We included 63 adults, of whom 27 were in the COVID-19+ group and 36 were in the COVID-19-group, of whom 24 were nonsusceptible. COVID-19+ and susceptible patients were older and had a higher prevalence of comorbidities than COVID-19- and nonsusceptible patients. The general characteristics of the study population are described in **Table 1**.

**Table 1.**
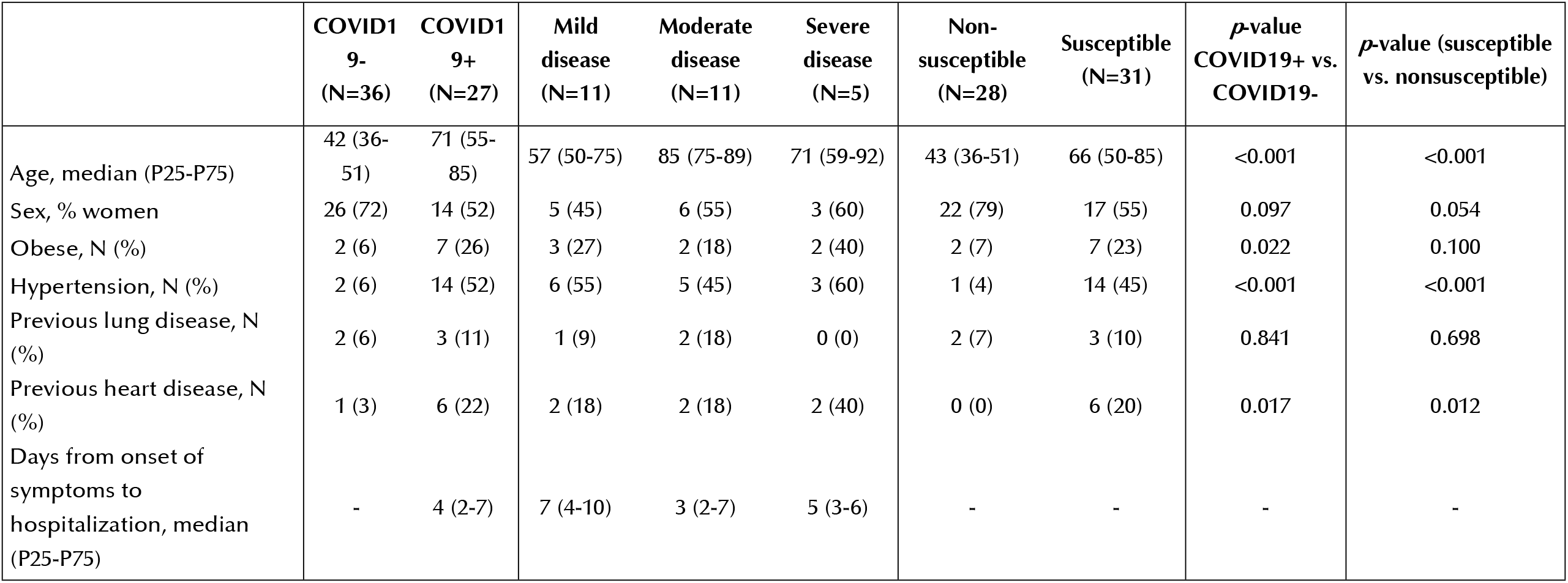
General characteristics of the study population.

### Differences in metabolic profiles according to COVID-19 disease status and SARS-CoV-2 susceptibility

Using an untargeted metabolomics approach and after data matrix filtration, 166 features (pairs of *m/z* –RT) were obtained in plasma samples with proper reproducibility. We first evaluated the differences between the metabolomes of COVID-19+ and COVID-19-participants, as well as between susceptible and nonsusceptible individuals, by building principal component analysis (PCA) and partial least squares-discriminant analysis (PLS-DA) score plots. As shown in **Fig S1 AB** and **Fig S2 AB**, the metabolomes of COVID-19+ vs. COVID-19-, as well as susceptible vs. nonsusceptible participants, drastically differed, indicating that the metabolomic fingerprints predicted the study group. Then, we performed orthogonal partial least squares-discriminant analysis (OPLS-DA) and found that the separation was totally explained through PC1 (**Fig 1**). The *p*-values for the OPLS-DA models were 2.10 × 10^−19^ and 4.20 × 10^−17^ for COVID-19 disease and COVID-19 susceptibility, respectively, corroborating previous observations. Using predefined statistical criteria for variable selection (VIP ≥ 1 and │p(corr)│ ≥ 0.5), we defined 10 metabolites predicting COVID-19 status and 11 predicting SARS-CoV-2 susceptibility (**Table S1**).

**Fig 1.**
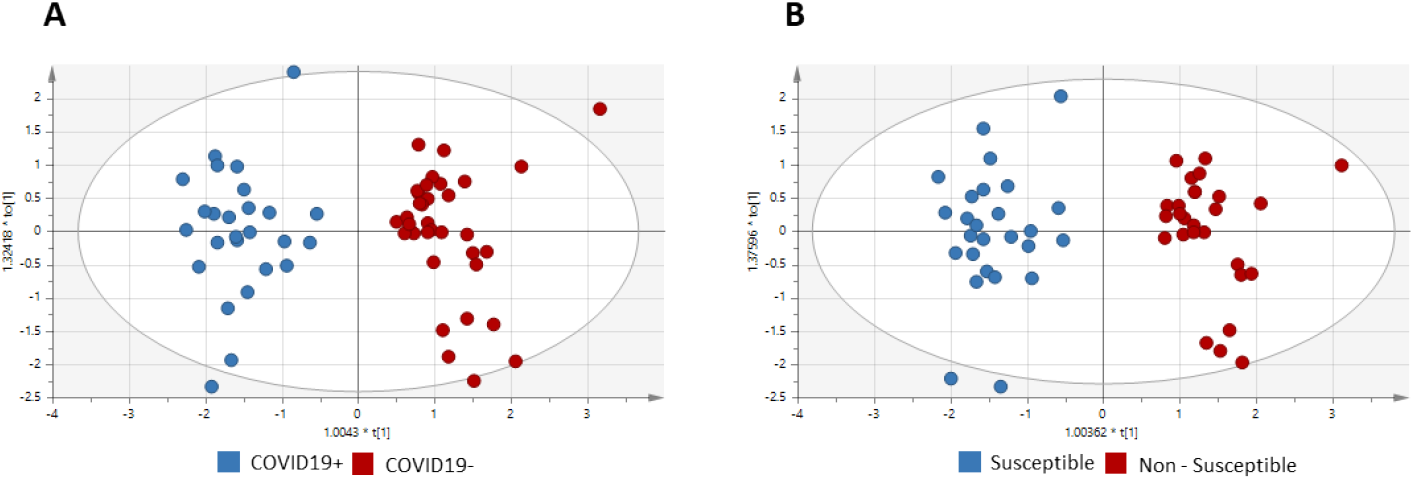
Untargeted metabolomic profiles of COVID19+ vs COVID19- and susceptible vs. non-susceptible participants using supervised OPLS-DA models for CE-MS data. (A) Plot A represents the comparison of COVI19+ and COVID19- individuals (R^2^ = 0.878, Q^2^ = 0.813), and CV-ANOVA (*p*-value = 2.10 × 10^−19)^. (B) Plot B represents the comparison of susceptible and non-susceptible participants with R^2^ = 0.902, Q^2^ = 0.817, and CV-ANOVA *p*-value = 4.20 × 10^−17^. Models were validated by permutation testing and CV-ANOVA (14,15). Hydroxychloroquine, initially found to be significant, was removed from all statistical analysis as it was empirically used to treat COVID19 at the time of sample collection.

### Metabolic profile differences associated with COVID-19 clinical severity

We then performed subgroup analyses separating COVID-19+ participants by clinical severity. While no differences in untargeted metabolomic profiles were found in the PCA (**Fig S1 C**), inspection of the PLS-DA score plots (**Fig S2 C**) showed clear clustering that did not meet the prespecified validation criteria. Pairwise comparisons of OPLS-DA models of all 3 categories fulfilled the validation criteria, indicating that there were statistically significant differences in the metabolomes of mild vs. severe and between moderate vs. severe cases (**Fig 2**). Similar to other COVID-19 and susceptibility studies, a total of 8 metabolites, including creatine, citrulline and 6 unknown features, were identified as predictors of greater disease severity (VIP ≥ 1 and │p(corr)│ ≥ 0.5) (**Table S1**).

**Fig 2.**
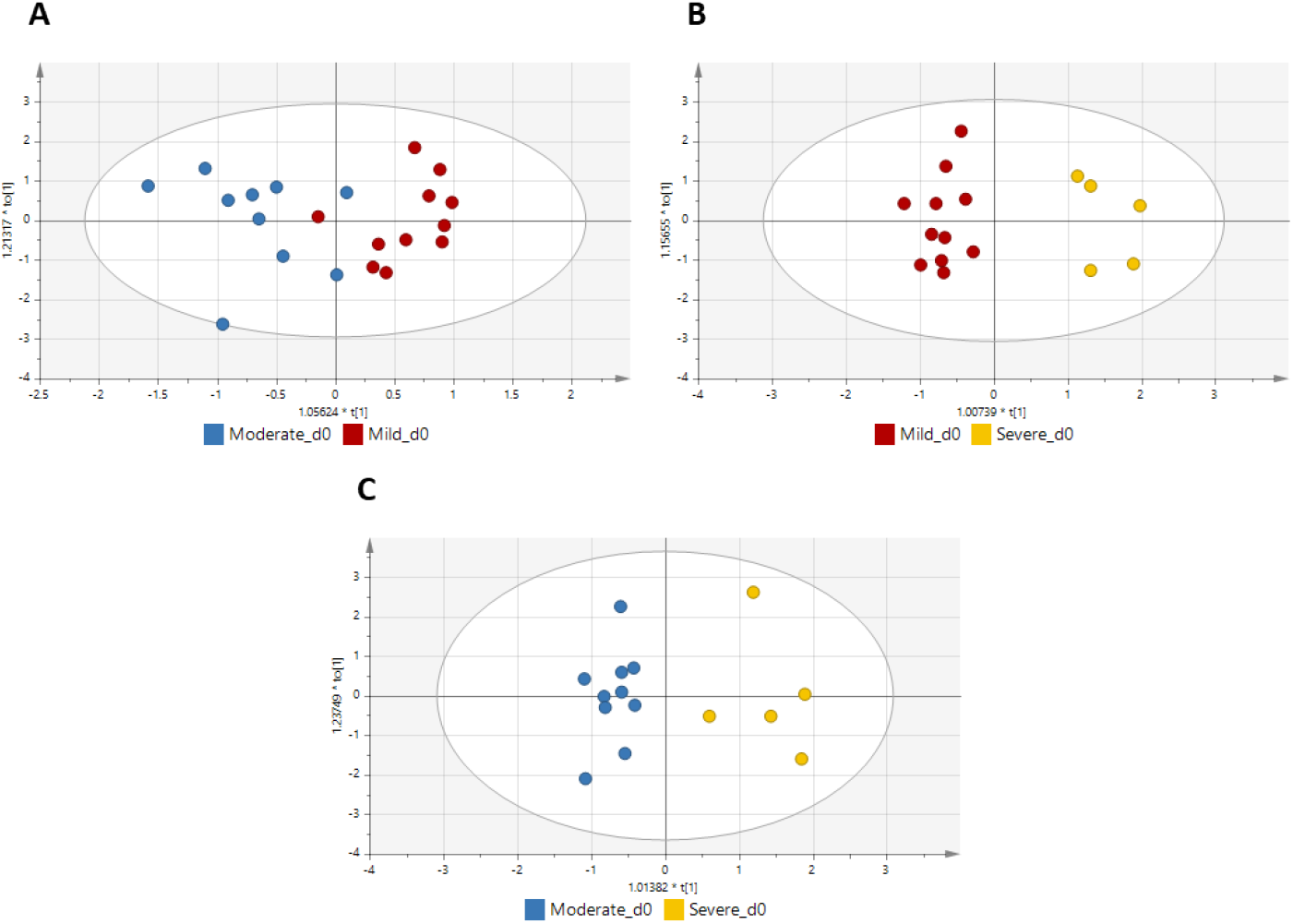
Untargeted metabolomic profiles of participants with COVID19 according to clinical severity using supervised OPLS-DA models for CE-MS data. (A) Mild vs. moderate disease; R^2^ = 0.713, Q^2^ = 0.009, and CV-ANOVA *p*-value = 0.997. (B) Mild vs. severe disease; R^2^ = 0.929, Q^2^ = 0.675, and CV-ANOVA *p*-value = 0.010. (C) Moderate vs. severe disease; R^2^ = 0.897, Q^2^ = 0.636, and CV-ANOVA *p*-value = 0.027.

### Longitudinal changes in the metabolomes of participants with COVID-19

We then sought to assess the effect of time on the metabolomes of participants with COVID-19 following a similar strategy. A clear separation between baseline and day 8 was found for mild and moderate cases (**Fig S1 D1-D3; Fig S2 D1-D2**). For severe cases, the PLS-DA model could not be fitted due to the limited availability of paired samples. Validated OPLS-DA models (**Fig 3**) showed that the longitudinal differences detected for mild and moderate cases were statistically significant (CV-ANOVA *p*-value < 0.05 and R^2^ - Q^2^ < 0.3). We found 10 metabolites whose abundance differed from baseline to day 8 in mild cases and 7 in moderate cases (VIP ≥ 1 and │p(corr)│ ≥ 0.5), see S1 Table).

**Fig 3.**
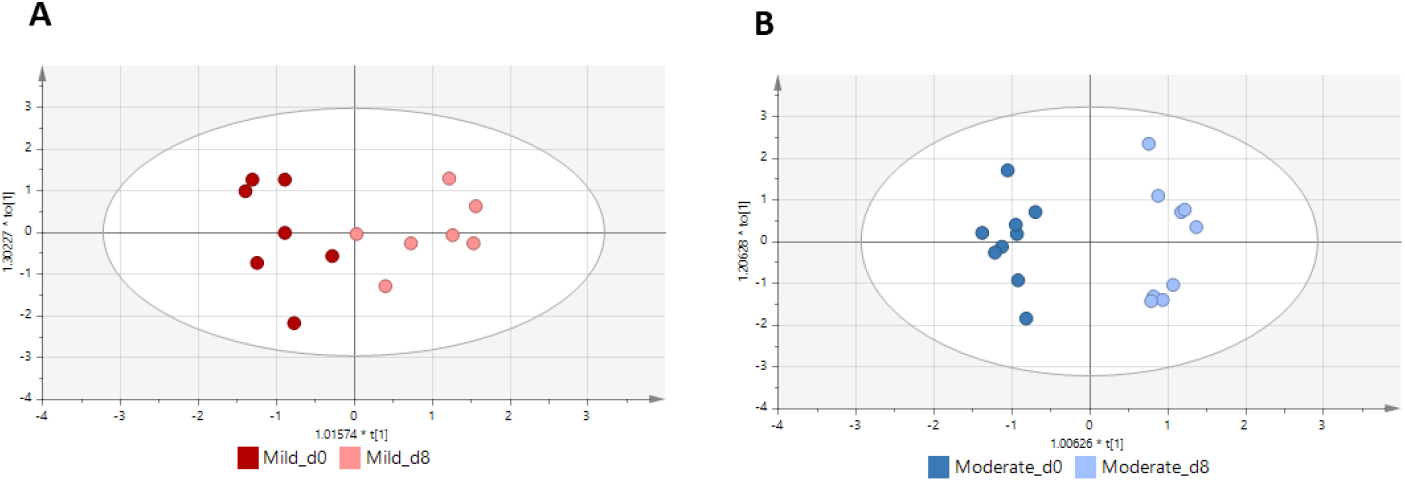
Untargeted metabolomic profiles at baseline and day 8 of participants with mild and moderate COVID19 supervised OPLS-DA models for CE-MS data. (A) Plot A represents the differences in mild cases (R^2^ = 0.816, Q^2^ = 0.596; CV-ANOVA *p*-value = 0.062). (B) Plot B represents the differences in moderate cases (R^2^ = 0.961, Q^2^ = 0.716; CV-ANOVA *p*-value = 0.014).

### Complementary characterization of metabolomic predictors of COVID-19 disease status and susceptibility

To visually summarize the metabolite fingerprint associated with COVID-19 disease and SARS-CoV-2 susceptibility, we represented the abundance of the metabolites identified by univariate analysis followed by multivariate statistical analysis as predictors of each condition in heatmaps with hierarchical clustering (**Fig 4, Fig S3 and Fig S4**). As shown in the heatmap (**Fig 4**), COVID-19+ (mild, moderate, and severe patients) had more similarities than COVID-19-individuals. The heatmap shows clear patterns of the distinct metabolic biomarkers for each group.

**Fig 4.**
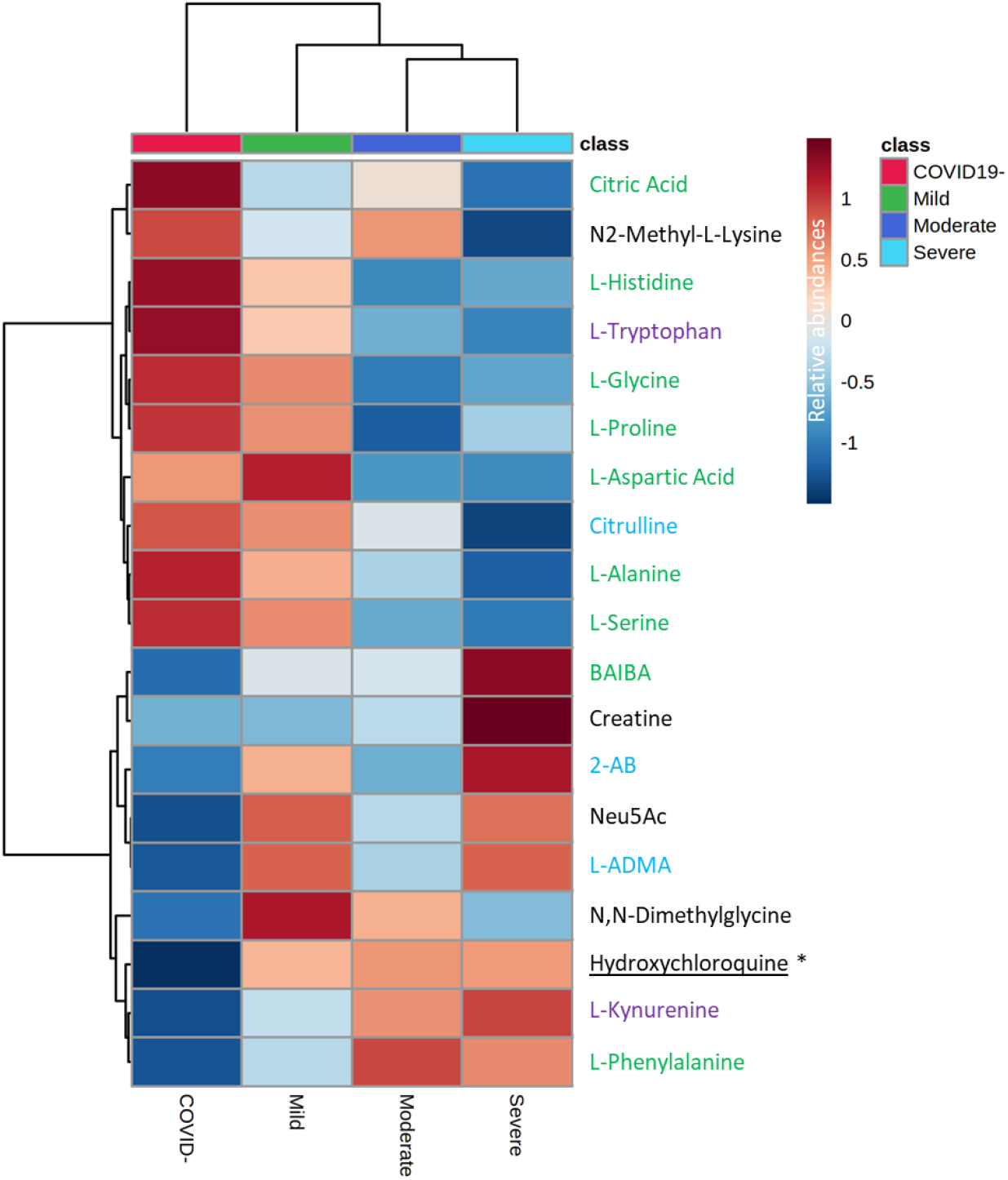
Heatmap with group average of statistically significant metabolites detected in human plasma samples by CE-MS modified by virus SARS-CoV-2 virus infection. In green, metabolites involved in TCA cycle. In purple, those involved in kynurenine pathway. In blue those compounds of the nitric oxide or are related with NO regulation.

ANOVA-simultaneous component analysis (ASCA) identified age as the only factor significantly associated with the outcome. Thus, we further assessed the metabolites previously identified as predictors of COVID-19 disease severity or susceptibility controlling for age using ANCOVA (Tables S2 and S3). Of them, NG, NG’-dimethyl-L-arginine (L-SDMA), L-cystine and L-carnitine lost statistical significance. L-Kynurenine and citric acid remained significantly predictive of COVID-19 disease and SARS-CoV-2 susceptibility, respectively. The selection of metabolites that could be fully characterized and their size effects are summarized in **Table 2**.

**Table 2.** Fold change of metabolite abundance in plasma samples associated with COVID19 disease status and susceptibility.

| Compound | COVID19+ vs. COVID19- | Susceptible vs. Nonsusceptible | Severe vs. Mild | COVID19+ day 8 vs baseline |
| --- | --- | --- | --- | --- |
| L-Glycine | nssd | ↓ 0.86 | nssd | nssd |
| L-Alanine | ↓ 0.85 | ↓ 0.85 | nssd | nssd |
| N,N-Dimethylglycine | nssd | nssd | nssd | ↑ 1.21 |
| 2-Aminobutyric acid | ↑ 1.37 | nssd | nssd | nssd |
| 3-Aminoisobutyric acid | ↑ 1.66 | nssd | nssd | nssd |
| L-Serine | ↓ 0.86 | ↓ 0.85 | nssd | nssd |
| L-Proline | ↓ 0.83 | ↓ 0.84 | nssd | nssd |
| Creatine | nssd | nssd | ↑ 2.47 | nssd |
| L-Aspartic acid | nssd | ↓ 0.87 | nssd | nssd |
| L-Histidine | ↓ 0.81 | ↓ 0.80 | nssd | nssd |
| N2-Methyl-L-Lysine | nssd | nssd | nssd | ↑ 1.5 |
| L-Phenylalanine | ↑ 1.16 | nssd | nssd | nssd |
| Citrulline | ↓ 0.81 | ↓ 0.80 | ↓ 0.56 | nssd |
| Citric acid | ↓ 0.80 | nssd | nssd | nssd |
| NG,NG-Dimethyl-L-Arginine | ↑ 1.17 | nssd | nssd | nssd |
| L-Tryptophan | ↓ 0.68 | ↓ 0.67 | nssd | nssd |
| L-Kynurenine | ↑ 1.53 | nssd | nssd | nssd |
| N-Acetylneuraminic acid | ↑ 1.81 | ↑ 1.8 | nssd | nssd |

Blue color denotes the fold change representing the increase of metabolite abundance and red color represents the decreases (see Table
S1 for additional information).

## Discussion

To our knowledge, this study is the first to evaluate the plasma metabolomic profile of individuals with severe COVID-19 and SARS-CoV-2 susceptibility by CE-MS. Our work demonstrates the potential of CE-MS to unveil new plasma biomarkers of COVID-19 and SARS-CoV-2 susceptibility and allows a deeper advancing of the metabolic consequences of SARS-CoV-2 infection (**Fig 5**).

**Fig 5.**
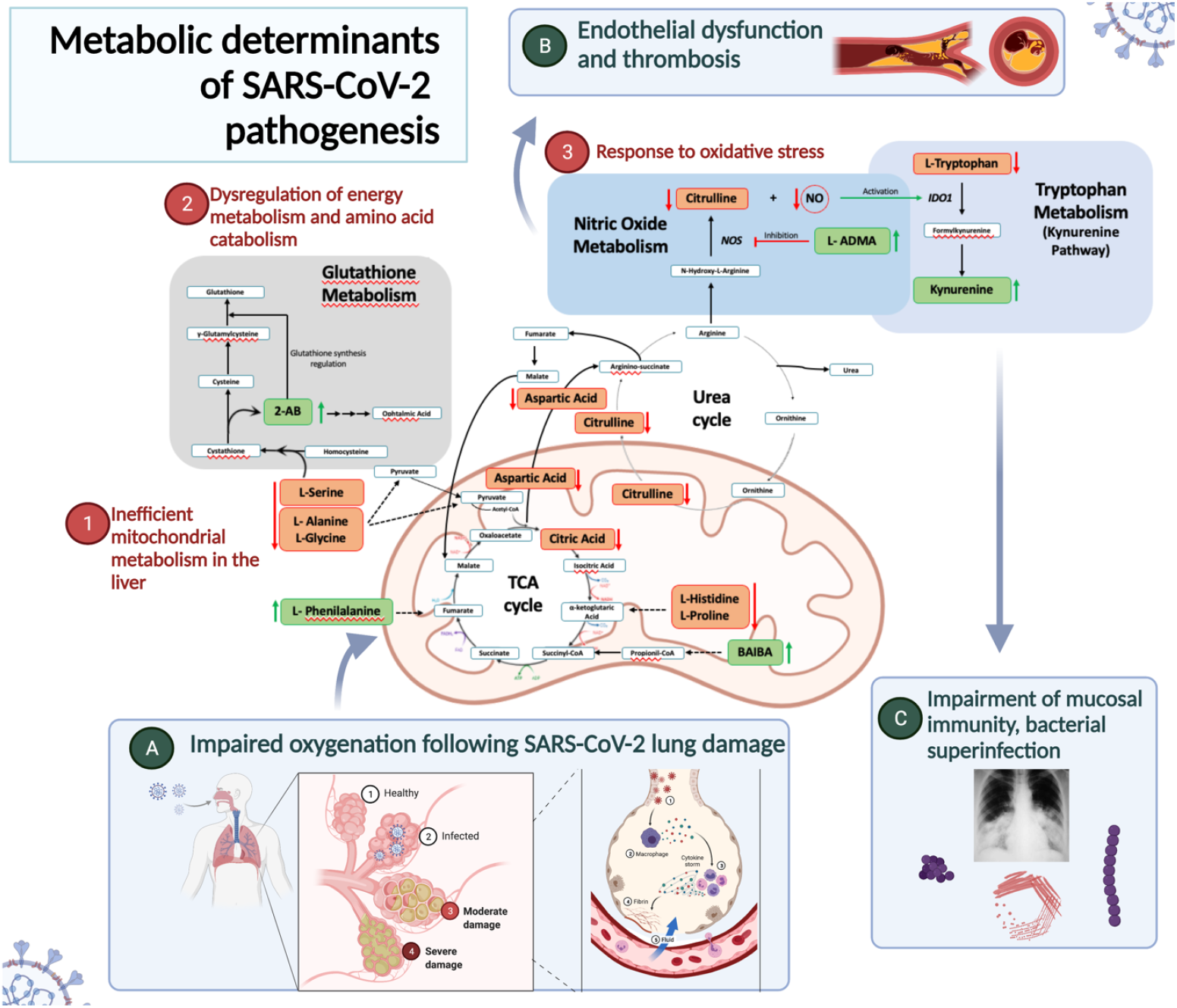
A model of the metabolic pathways implicated in COVID19 pathogenesis. Impairment of blood oxygenation following SARS-CoV-2 damage results in 1) inefficient mitochondrial metabolism in the liver, resulting in dysregulation of the urea cycle citrulline decreases, phenylalanine increases; 2) dysregulation of energy metabolism and amino acid metabolism, resulting in decreased L-serine, L-alanine, and L-serine; 3) activation of oxidative stress response, resulting in BAIBA accumulation, L-ADMA upregulation, and induction of the kynurenine pathway, which impairs mucosal immunity, allowing bacterial superinfections. Figure generated using biorender.com.

Among the significant metabolites, we found that the citrulline concentration decreases over the course of COVID-19 disease, but low levels early on in the course of the disease are associated with greater clinical severity. This finding is consistent with those reported in a recent work, where carbamoyl phosphate levels, a substrate for citrulline biosynthesis in the mitochondria of liver cells, decreased with greater disease severity (1). Because citrulline is an intermediate in the urea cycle and a byproduct of the enzymatic production of nitric oxide from arginine (16), these findings point to either dysregulation in the urea cycle or liver dysfunction as the underlying mechanism explaining the links between this metabolite and COVID-19. Furthermore, increased levels of circulating phenylalanine, which were found to be associated with COVID-19 in our study, have also been reported in patients with hepatic fibrosis, acute hepatic failure and hepatic encephalopathy as well as in COVID-19 disease (5).

Apart from phenylalanine, other amino acids (AAs) were found to be significantly different between the groups (Table 2). Among them, L-glycine, L-alanine, L-serine, L-proline, L-aspartic acid and L-histidine were downregulated in patients. Previous studies have revealed that SARS-CoV-2 infection dysregulates pathways linked to energy production and amino acid catabolism (17,18). In a murine model of SARS-CoV-2, Li *et al*. found several genes commonly downregulated in multiple organs that led to significant enrichment in pathways related to oxidative phosphorylation and the electron transport chain (17). As the tricarboxylic acid (TCA) cycle is connected to the electron transport chain, they also analyzed genes associated with the TCA cycle. They found that several TCA cycle genes were downregulated and that TCA cycle metabolites were decreased in animal serum (17).

Apart from the AAs that lead to intermediates of the TCA cycle that were downregulated in the COVID-19+ group, the significant downregulation of citrate also suggested that SARS-CoV-2 results in inefficient mitochondrial metabolism (18,19), which can be interpreted as the metabolic response to impaired oxygenation secondary to lung damage (9). Citrate is a direct TCA cycle metabolite obtained by the action of citrate synthase from oxaloacetate. The gene encoding this enzyme exhibits decreased expression (17). Different genes, proteins and/or metabolites involved in the TCA cycle have been found to be suppressed or downregulated in individuals with COVID-19 (18,19).

An intriguing finding in our study is the upregulation of 3-aminoisobutyric acid (BAIBA) associated with COVID-19. BAIBA is a catabolite of thymine and valine metabolism that has been proposed as a novel regulator of carbohydrate and lipid metabolism associated with aerobic exercise (20). Although little is known about the implications of BAIBA in pathogenesis, the fact that two enantiomers of BAIBA (R-BAIBA and S-BAIBA) are ultimately metabolized in mitochondria further supports the idea that mitochondrial and TCA cycle abnormalities are a metabolic hallmark of COVID-19 pathogenesis, as also indicated by the abnormalities detected in amino acid and citrate metabolism (21). As BAIBA is primarily metabolized by mitochondria, the accumulation of BAIBA in patients with COVID-19 could be explained by a reduction in mitochondrial functionality and TCA cycle suppression following impairment of blood oxygenation. To our knowledge, BAIBA has never been proposed as a putative metabolite involved in COVID-19 disease. This result is of special interest not only to further investigate BAIBA as a novel biomarker for COVID-19 disease but also to elucidate its role in metabolism under physiological stress conditions or hypoxemia.

We also found evidence that SARS-CoV-2 affects metabolic pathways implicated in endothelial dysfunction, thrombosis, and cardiovascular disease. First, nitric oxide synthase (NOS) is an enzyme that catalyzes the production of citrulline and nitric oxide (NO) from arginine. This enzyme is inhibited by asymmetric dimethylarginine (L-ADMA), which is upregulated in COVID-19 patients and is an endogenous competitor of arginine, the nitric oxide precursor (22). L-ADMA has been associated with elevated oxidative stress (23). The higher L-ADMA concentrations found in individuals with COVID-19 suggest inhibition of NOS activity, which would ultimately result in decreased levels of NO. Because NO is among the principal redox molecules exploited by the immune system as a defensive mechanism, NO has been implicated in the control of viral replication, including that of HIV, influenza A and B, and vaccinia virus (24,25). Because it is as-yet unexplained how SARS-CoV-2 produces severe endothelial injury, widespread thrombosis and microangiopathy (26), our findings offer a new mechanistic explanation for this hallmark of SARS-CoV-2 pathogenesis and point to the nitric oxide synthesis pathway as a potential therapeutic target. Second, 2-aminobutyric acid (2-AB) and N-acetylneuraminic acid (Neu5Ac) were also upregulated in the COVID-19+ group. 2-AB is a marker that seems to be a compensatory mechanism to oxidative stress (27) and has been implicated in the modulation of glutathione metabolism in the myocardium (28). This finding indicates that 2-AB deserves further attention as a biomarker of the myocardial dysfunction associated with COVID-19 (29). Finally, Neu5Ac is the most widespread form of sialic acids and is a family of compounds with a broad range of implications in human physiology (30). Because Neu5Ac concentrations have been correlated with the development of cardiovascular disease via RhoA signaling pathway activation (31,32), the fact that we found higher Neu5Ac concentrations associated with COVID-19 provides a new pathway possibly linked to the excess risk of cardiovascular diseases associated with SARS-CoV-2.

Inflammation gained early attention as a crucial mechanism of SARS-CoV-2 pathogenesis (33). Indoleamine-2,3-dioxygenase-1 (IDO1), which is involved in tryptophan catabolism via the kynurenine pathway, is correlated with epithelial barrier disruption, bacterial translocation and inflammation in other viral infections (34). Induction of IDO1 results in the production of kynurenine derivatives with immunosuppressive effects, impairing mucosal immunity and promoting bacterial translocation and higher mortality (35). Impairment of the kynurenine pathway, resulting in reduced tryptophan (Trp) and elevated kynurenine (Kyn) levels associated with COVID-19, has previously been reported (3,7,36). Our data reveal not only the same tendency for Trp and Kyn but also the increasing tendency of the Kyn/Trp ratio with severity. This ratio has previously been associated with renal insufficiency in patients with SARS-CoV-2 and in many other diseases, such as inflammatory lung disease (5,37). Strikingly, IDO activity is induced by interferon-gamma (IFN-γ), as well as other cytokines and mediators (38,39), and it is inhibited in oxidative stress conditions by NO (39,40). Considering the reduction in NO synthesis mentioned previously, the alterations observed in the kynurenine pathway could be a result of the aforementioned metabolic abnormalities and result in further impairment of mucosal immunity, providing an explanation for the significant rates of bacterial pneumonia associated with COVID-19 (35).

The major strengths of our study include 1) the inclusion of COVID-19 cases in an early phase since the onset of symptoms, 2) the assessment of a special population of nonsusceptible individuals, 3) the high-throughput CE-MS method used to characterize the metabolome of the study participants, and 4) the inclusion of follow-up samples to assess the longitudinal variations of the plasma metabolites in a subset of participants. Our study is also subject to some limitations. First, the samples were collected during the first COVID-19 wave in Madrid. It is unknown yet whether the emerging SARS-CoV-2 variants could lead to different metabolic consequences. Second, as expected, cases in the severe group were older and had more comorbidities than milder cases, so we considered potential confounders in our statistical approach. Third, in the subgroup analyses separated by clinical severity, the statistical power to detect differences in metabolite abundances was lower due to the smaller sample sizes.

In summary, in this work examining for the first time the metabolic changes associated with COVID-19 by CE-MS, we report the discovery of new plasma biomarkers for COVID-19 that provide mechanistic explanations for the clinical consequences of SARS-CoV-2, including mitochondrial and liver dysfunction as a consequence of hypoxemia (citrulline, citrate and BAIBA), energy production and amino acid catabolism (L-glycine, L-alanine, L-serine, L-proline, L-aspartic acid and L-histidine), and endothelial dysfunction and thrombosis (citrulline, L-ADMA, 2-AB, and Neu5Ac), and we found interconnections between these pathways (**Figure 5**). These biomarkers deserve further attention as biomarkers of SARS-CoV-2 susceptibility and COVID-19 clinical severity and as potential targets for interventions.

## Material and methods

### Reagents

All reagents, solvents and standards used for sample treatment and subsequent analysis are described in the Supporting Information.

### Patient enrollment and sample collection

We analyzed data from adults recruited at Hospital Universitario Ramón y Cajal, Madrid, Spain. Participants had confirmed SARS-CoV-2 (COVID-19+ group) infection by PCR from nasopharyngeal swabs, sputum, or lower respiratory tract secretions within the first 7 days from the onset of symptoms and were classified according to clinical severity as follows: mild disease, defined as those without a need for supplemental oxygen and who were asymptomatic one week after diagnosis; moderate disease, defined as the presence of bilateral radiologic infiltrates or opacities and clinical assessment requiring supplemental oxygen; and severe disease, defined as the development of acute respiratory distress syndrome (41). Hospitalized participants provided samples at baseline and 8 days later. Participants without SARS-CoV-2 (COVID-19-group) were asymptomatic subjects with a negative PCR from nasopharyngeal swabs. We considered adults to be “susceptible” when they had positive IgG for SARS-CoV-2 or previous COVID-19 confirmed by polymerase chain reaction (PCR) from nasopharyngeal exudate. Nonsusceptible adults were healthy healthcare workers who had been on duty for at least three months in COVID-19 wards or intensive care units and reported at least three high-risk exposures to SARS-CoV-2 (42) without having experienced symptoms suggestive of SARS-CoV-2 infection, were persistently negative for SARS-CoV-2 PCR testing and did not have SARS-CoV-2 IgM and IgG in plasma. The most frequent exposure was largely unprotected exposure to aerosol-generating procedures or patient secretions and close contact without face masks with other confirmed cases of COVID-19. We measured SARS-CoV-2 antibodies by indirect chemiluminescence immunoassay (Vircell, Granada, Spain).

Cryopreserved plasma was processed for virus inactivation by adding 1500 µL of cold methanol:ethanol (MeOH:EtOH) in a 1:1 (v/v) proportion to 500 µL of plasma. Then, samples were vortex-mixed for 1 min, incubated on ice for 5 min and centrifuged at 16,000 x *g* for 20 min at 4 °C to precipitate and remove proteins. The clean upper layer or supernatant, which contained the metabolites of interest, was transferred to Eppendorf tubes and stored at -80 °C until analysis.

### Sample treatment

Two hundred microliters of frozen supernatant was thawed on ice and evaporated to dryness using a SpeedVac Concentrator System (Thermo Fisher Scientific, Waltham, MA). Then, it was resuspended in 100 µL of 0.2 mM methionine sulfone (MetS) in 0.1 M formic acid. Samples were vortex-mixed for 1 min, transferred to a Millipore filter (30 kDa protein cutoff) and centrifuged for 40 min at 2000 x*g* at 4 °C. Finally, the ultrafiltrate was transferred to a CE-MS vial for analysis. Quality control samples (QC) were prepared by pooling equal volumes of plasma supernatant from each sample and were treated as previously described. Finally, blank solutions were also prepared with MeOH:EtOH (1:1, v/v).

### Nontargeted metabolomics by CE-MS

The plasma metabolome was analyzed by using a 7100 capillary electrophoresis (CE) system coupled to a 6230 time-of-flight mass spectrometer (TOF-MS) from Agilent Technologies equipped with an electrospray ionization (ESI) source. The analysis was performed using a previously developed method (43) with the analytical conditions described in detail in the Supporting Information. The prepared QCs were analyzed at the beginning of the run to condition the CE system and then every seven randomized samples to reduce any time-related effect. The QCs were used not only to assess the reproducibility, stability and performance of the system but also to correct any signal deviation within the analytical sequence. A pair of blanks were injected at the beginning and end of the run to remove metabolites coming from the extraction solvent.

### Data processing

CE-MS raw data were checked using MassHunter Qualitative software (version 10.0) to determine the data quality, the system mass accuracy and the reproducibility of the QC sample and IS injections. Then, raw data were aligned and processed with MassHunter Profinder software (version 10.0 SP1). Molecular feature extraction (MFE) and batch recursive feature extraction (RFE) algorithms, both included in MassHunter Profinder software, were used to obtain the list of mass-to-charge ratios (*m/z*) and their corresponding abundances (43). The resulting list was imported in Microsoft Excel, and the data matrix was filtered before statistical analysis by removing metabolites with a percentage of coefficient of variation (% CV) greater than 30% in the QC samples. All the data processing steps are described in detail in the Supporting Information.

### Statistics

Multivariate (MVDA) and univariate (UVDA) statistical analyses were carried out to determine differences among groups. Different comparisons were performed to evaluate COVID-19 disease, disease severity, disease progression, and susceptibility. For this purpose, samples were labeled based on the comparison as infected or noninfected for disease diagnosis; susceptible or nonsusceptible for disease susceptibility; mild, moderate or severe at day 0 (d0) for disease severity; or day 0 and day 8 for disease progression. Then, the filtered matrix obtained in the previous step was processed by SIMCA-P version 15.0.2 (Umetrics, Umea, Sweden), MATLAB software (The MathWorks, Maticks, MA, USA), MetaboAnalyst 5.0 and SPSS version 24 (IBM SPSS Statistics) for different purposes. When needed, the intensity drop was corrected with the QC correction function included in the toolbox freely available online at https://github.com/Biospec/cluster-toolbox-v2.0. Statistical analysis is described in more detail in the Supporting Information. Briefly, unsupervised PCA was performed to visualize tendencies, determine the presence of outliers, and assess data quality by the explained variance (R^2^) and the predicted variance (Q^2^), considering as an appropriate value a difference between them of lower than 0.3 (15). Then, the supervised methods PLS-DA and OPLS-DA were performed followed by model validation. In those validated OPLS-DA models, variable selection was performed by using a variable influence on projection (VIP) and absolute value of p(corr) greater than 1.0 and 0.5, respectively (14). Afterwards, UVDA was performed simultaneously to assess the significance of each metabolite separately. In short, nonparametric tests were applied for the comparisons previously mentioned as follows: a) the Kruskal-Wallis test for disease severity (mild, moderate, and severe patients at d0) followed by a multiple comparison test; b) the Wilcoxon signed-rank test for disease progression; and c) the Mann-Whitney U test for COVID-19 disease and susceptibility. In all cases, the *p*-value had to be less than 0.05, and the false discovery rate at a level of α = 0.05 was controlled by the Benjamini-Hochberg correction test. Finally, ASCA was applied to study the influence associated with sex and age (44). When the ASCA model was not validated by permutation testing, analysis of covariance (ANCOVA) was carried out to eliminate the variability associated with age, sex or both (45).

### Metabolite identification

The selected features in the statistical step by UVDA or MVDA were tentatively identified based on the *m/z* of the metabolites and the relative mobility time (RMT) (RT_metabolite_/RT_MetS_) by using the CEU Mass Mediator (http://ceumass.eps.uspceu.es/mediator) (46), which is an ‘in-house’ useful tool for identification. This tool joins several databases, which are available online, such as METLIN (47), LIPIDMAPS (48), and KEGG (49), making the identification task faster and easier. Features assigned to metabolites have to fulfill an appropriate mass accuracy (maximum error mass of 15 ppm), as well as a comparable isotopic pattern distribution. Once metabolites were identified, confirmation was performed by injecting commercial standards, samples, and samples spiked with standards. Finally, for fragmentation pattern recognition, the QC sample was analyzed under the same analytical conditions as used in the previous analysis but applying different voltages in the MS fragmentor (150, 175 and 200 V) (50). It is important to point out that any drug associated with COVID-19 treatments that was identified among the significant metabolites was excluded from both MVDA and UVDA statistical analysis.

### Study approval

The study was carried out at the Ramón and Cajal University Hospital in Madrid (Spain) and was approved by the local Research Ethics Committee (,approval number 095/20). All subject unable to provide informed consent or witnessed oral consent with written consents by a representative were excluded.

## Contributors

O.E.A., S.S-V., and C.B study design and conceptualization; S.S-V., D.J., M.S-C., P.V., R.R., S.H., J.M-S. and S.M. recruited and clinical follow-up; D.J. handling of clinical specimens and data mining; O.E.A., C.B., measurement of plasma metabolites; O.E.A. and S.S-V statistical analysis; O.E.A. and C.B bioinformatic analyses; O.E.A. writing of the first version of the manuscript. All the authors reviewed and approved the manuscript.

## Acknowledgments

We thank to all patients and healthcare workers who participated in the study.

## Data availability

Participant’s metadata and abundances of the key metabolites are displayed in the supplemental table as a supplementary data file.

